# Specific oncogene activation of the cell of origin in mucosal melanoma

**DOI:** 10.1101/2024.04.22.590595

**Authors:** Swathy Babu, Jiajia Chen, Emily Robitschek, Chloé S. Baron, Alicia McConnell, Constance Wu, Aikaterini Dedeilia, Moshe Sade-Feldman, Rodsy Modhurima, Michael P. Manos, Kevin Y. Chen, Anna M. Cox, Calvin G. Ludwig, Jiekun Yang, Manolis Kellis, Elizabeth I. Buchbinder, Nir Hacohen, Genevieve M. Boland, Brian J. Abraham, David Liu, Leonard I. Zon, Megan L. Insco

## Abstract

Mucosal melanoma (MM) is a deadly cancer derived from mucosal melanocytes. To test the consequences of MM genetics, we developed a zebrafish model in which all melanocytes experienced CCND1 expression and loss of PTEN and TP53. Surprisingly, melanoma only developed from melanocytes lining internal organs, analogous to the location of patient MM. We found that zebrafish MMs had a unique chromatin landscape from cutaneous melanoma. Internal melanocytes could be labeled using a MM-specific transcriptional enhancer. Normal zebrafish internal melanocytes shared a gene expression signature with MMs. Patient and zebrafish MMs have increased migratory neural crest gene and decreased antigen presentation gene expression, consistent with the increased metastatic behavior and decreased immunotherapy sensitivity of MM. Our work suggests the cell state of the originating melanocyte influences the behavior of derived melanomas. Our animal model phenotypically and transcriptionally mimics patient tumors, allowing this model to be used for MM therapeutic discovery.

## Main Text

Mucosal melanoma (MM) is a cancer of pigment-producing melanocytes that arises in mucosal surfaces and has a poor prognosis with a 5-year survival rate of 27%, largely due to its increased metastatic behavior as compared to stage-matched cutaneous melanomas (CMs)^1^. MMs typically occur in sun-protected sites and thus have a lower UV mutational burden, which is thought to contribute to the low patient responses to immune checkpoint blockade^2^. In contrast, MMs have abundant large-scale genomic rearrangements, which result in oncogene amplifications and tumor suppressor deletions^3, 4, 5, 6^, whose molecular, phenotypic, and therapeutic consequences remain mysterious. Genomic studies have revealed that MM genomes rarely harbor activating BRAF^V600E^ mutations, making them ineligible for BRAF inhibitors. Activating KIT mutations are enriched in MM (15%) as compared to CM (2%), however responses to KIT small molecule inhibitors have been disappointing^4, 6, 7^. The discovery of therapeutics for MM will also likely benefit a subset of CM patients as demonstrated by the approval of tebentafusp for uveal melanoma, another rare subtype^8, 9^.

Though mutational and certain phenotypic differences between MM and CM have been identified, MM basic biology and therapeutic discoveries have been hampered by the lack of an animal model. Rapid genetic tumor modeling in zebrafish has contributed to major discoveries in CM biology^10, 11, 12^, the functional role of CRKL in acral melanoma^13^, as well as the role of SPRED1 deletion in KIT inhibitor resistance in MM^5^. Studies of spontaneously occurring MM in canines have been fruitful^14^, but there remains a need to develop genetically engineered animal models to determine functional drivers of MM in patients.

By modeling a combination of genetic changes that occur in MM patients including CCND1 amplification and PTEN and TP53 deletion, we have developed the first *bona fide* genetically engineered animal model of human MM. These CCND1-amp; *pten a/b*-/-; *tp53*-/- fish develop melanoma only *inside* the animal. Zebrafish MMs have a distinct epigenetic state compared to CM. A MM-specific transcriptional enhancer specifically labels a normal internal zebrafish melanocyte population, nominating a cell of origin. Single-cell RNA sequencing of adult internal and cutaneous melanocytes identifies *internal* melanocytes as the cell of origin for zebrafish MMs. Patient MMs share several key gene expression signatures with the zebrafish model including increased migratory neural crest expression profiles and decreased antigen presentation. Our data are consistent with MM originating from a unique melanocyte population whose epigenetic cell state contributes to lethality of this disease.

## Results

### Zebrafish model recapitulates localization of human MM

*CCND1* and *CDK4* are among the most significantly amplified genes in MM^3, 4, 5, 6^. *CCND1* is amplified in a mutually exclusive pattern with *CDK4*^3^ with amplification of either *CCND1* or *CDK4* present in 40% of MMs (27/63)^6^ as compared to 9.6% of CM patients (n=35/363)^15^. Amplifications of either *CCND1* or *CDK4* occur at a significantly higher frequency in tumors that are wild type for BRAF or NRAS, suggesting that CCND1 could be an independent melanoma driver^3^.

To test whether CCND1 could be an independent melanoma driver, we utilized the MAZERATI^5^ zebrafish rapid modeling system, which allows melanocyte-specific expression or loss-of-function of any gene. The MAZERATI system was utilized in zebrafish that lack *mitfa,* the master transcriptional regulator of melanocyte development, and thereby lack melanocytes. Melanocytes are rescued with vectors that express Mitfa and in a cell-autonomous melanocyte-specific manner, express or CRISPR-delete genes of interest under the control of an exogenous *mitfa* promoter. Transparent *mitfa^-/-^; roy^-/-^*fish (hereafter referred to as Casper zebrafish) were utilized so that the location of melanoma onset could be visually discerned. Using the MAZERATI system in Casper zebrafish, we engineered melanocyte-specific expression of human *BRAF^V600E^* and CRISPR-deletion of *tp53,* which led to cutaneous melanomas as expected^5^ (Fig. 1A, top). Melanocyte-specific expression of *CCND1* with CRISPR-deletion of both *pten a/b* and *tp53* led to pigmented tumors consistent with melanoma, however these tumors only arose *inside* the fish, which we have not previously observed (n=22/205) (Fig. 1A, bottom; Fig. 1B). These fish also developed tumors that were pigment free (95/205), but these were not further studied. Rare fin (fish acral) and eye (fish uveal) melanomas were also observed. This was the first time we observed melanoma being driven in the MAZERATI model in the absence of an activating MAPK pathway driver (e.g. BRAF^V600^ or NRAS^Q61^). To determine the rate that this combination of drivers occurs in patients, we analyzed targeted sequencing data from 57 MM samples previously published ^16^ in addition to 27 newly sequenced patients. Out of 86 MM, 45% of patients had deleterious alterations in PTEN (39/86), 36% in TP53 (31/86), 12 had amplifications in CCND1, and three patients had all three (Fig. S1A, Data S1). These data show that amplification of CCND1 and loss of PTEN and TP53 occur in MM patients. Together these data indicate that a population of internal melanocytes in zebrafish is susceptible to melanoma transformation upon melanocyte-specific expression of *CCND1* and loss of *pten a/b* and *tp53*.

**Figure 1:**
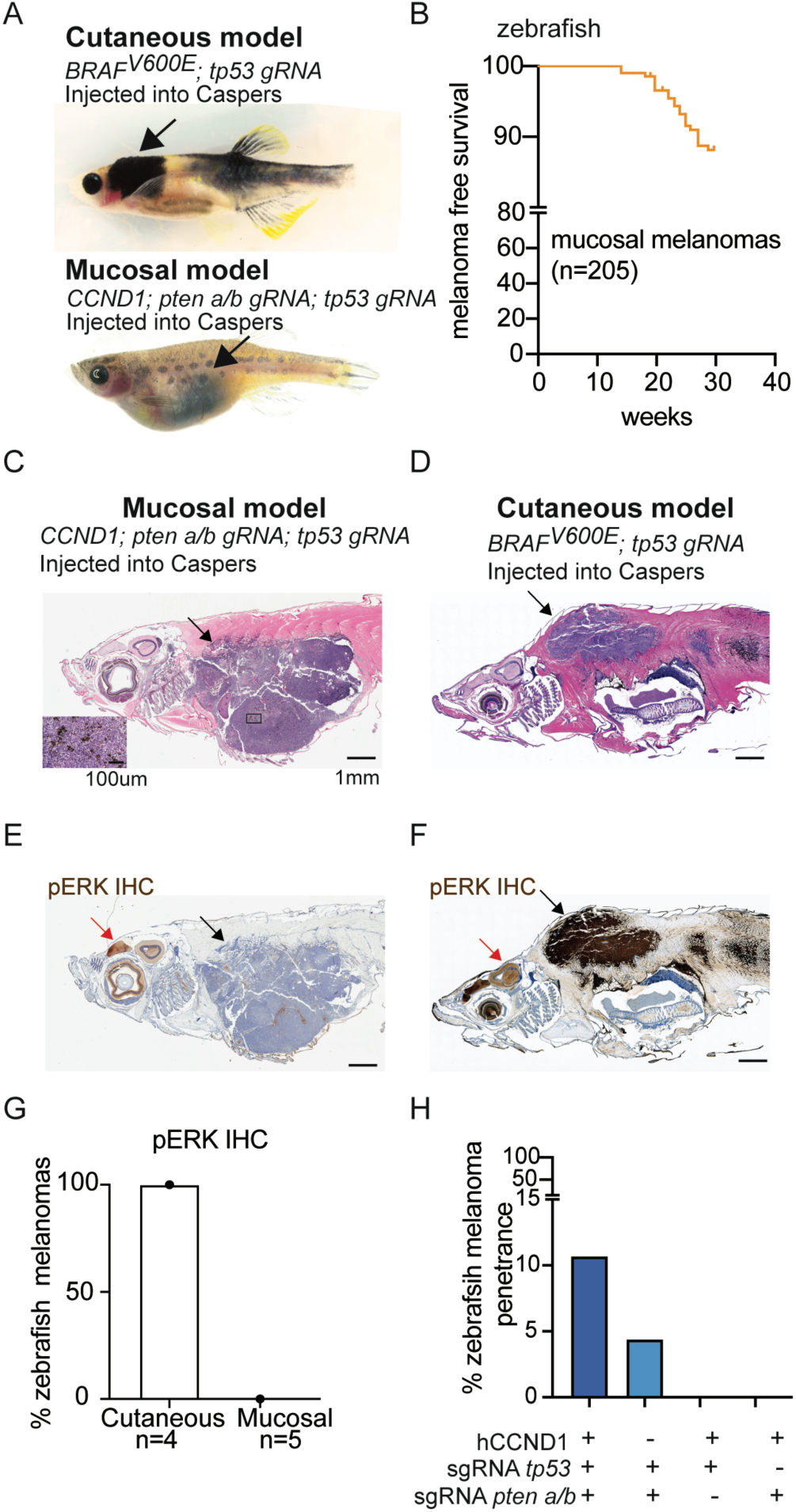
Zebrafish model recapitulates localization of human MM. A) Light micrographs of zebrafish melanoma generated from melanocyte-specific expression or CRISPR-deletion in transparent “Casper” zebrafish. Cutaneous model (upper). Mucosal model (lower). B) MM free survival of zebrafish with melanocyte-specific expression of human CCND1 and melanocyte-specific CRISPR of *pten a* and *b* and *tp53.* C-F) Zebrafish melanoma histology. C-D) H&E stain of MM (C) and CM (D). E-F) phospho ERK (pERK) immunohistochemistry (IHC) of MM (E) and CM (F). Scale bar = 1mm; inset scale bar = 100um. Black arrow = tumor, red arrow = brain. G) % zebrafish melanomas pERK positive. H) % zebrafish MM penetrance.

Histology of the engineered internal tumors demonstrated that the dominant mass was located in or just anterior to the abdominal cavity, indicating that the tumor likely initiated from melanocytes that normally reside near this location (Fig. 1C vs. 1D). Histologic examination demonstrated that these tumors contained pigment, supporting that they are melanomas (Fig. 1C, inset). Next generation sequencing of PCR products that span the CRISPR cut sites of *ptena, ptenb* and *tp53* demonstrated that all three genes were targeted, and every detected allele was out-of-frame (predicted to cause loss-of-function) as is expected with tumor suppressors (Fig. S1B-C). CCND1 immunohistochemistry (IHC) confirmed that all MMs checked expressed human CCND1 (n=5) while the CMs lacked CCND1 (n=4) (Fig. S1D-F). As the MAPK pathway is almost universally activated in CMs, and the zebrafish internal melanomas lack an obvious MAPK pathway driver mutation, we asked whether CCND1 expression activates MAPK. IHC for pERK, a strong and specific marker of MAPK signaling, showed that all CMs (n=4) and no MMs (n=5) were positive for pERK staining (Fig. 1E vs. 1F, quantification in Fig. 1G), while control pERK staining was observed in the brains of all animals (red arrow, Fig. 1E and F). These data demonstrate that CCND1 expression along with loss of *pten a/b* and *tp53* is sufficient to cause internal melanoma in zebrafish.

To determine which genetic changes were required, we took a ‘minus 1’ approach for each of the three drivers using the MAZERATI model in Casper zebrafish. Expression of CCND1 with CRISPR-deletion of *pten* a/b or *tp53* did not lead to internal melanoma development, while CRISPR-deletion of *pten* a/b and *tp53* without CCND1 expression resulted in a small number of internal melanomas (Table 1, Fig 1H). These data indicate that both *pten a/b* and *tp53* loss are required for the onset of internal melanomas, while CCND1 expression amplifies the penetrance of *ptena/b* and *tp53* loss. These data show that melanocytes inside the fish, analogous to human mucosal melanocytes, are susceptible to melanoma transformation in the absence of a strong MAPK driver mutation. These data suggest that this model represents a genetically engineered model of human MM and that MM derives from a conserved cell of origin distinct from that of CM.

**Table 1:** Zebrafish MM penetrance.

| Genotype | Melanoma |
| --- | --- |
| CCND1; pten a/b sgRNA; tp53 sgRNA | 11% (22/205) (internal only) |
| CCND1; tp53 sgRNA | 0% (0/100) |
| CCND1; pten a/b sgRNA | 0% (0/100) |
| tp53 sgRNA; pten a/b sgRNA | 5.6% (5/89) (internal only) |

### Zebrafish MM has a distinct cellular state from CM

To determine whether zebrafish MM and CM have distinct gene expression patterns, we sequenced polyA-selected bulk RNA from zebrafish MM (n=5) and CM (n=3). 742 genes were significantly differentially expressed between MM and CM, with 320 genes upregulated and 422 genes downregulated in MM (Fig. 2A). *sox10* and *mitfa* were both highly expressed and not significantly different between MM and CM, confirming that these tumors are melanomas (*sox10,* q= 0.68; *mitfa,* q=0.24), although both were expressed at higher levels in the MM model (Fig. S2A). In the MMs, the most significant downregulated pathway by GO-term analysis was “negative regulation of MAPK activity”, including many direct targets of MAPK signaling. Less activation of MAPK target genes was expected since these animals lack an activating MAPK driver mutation. Surprisingly, *tfap2a,* a gene typically highly expressed in both patient and zebrafish CMs^17^ was significantly downregulated in the internal melanomas (Fig 2A). Instead the ortholog, *tfap2b,* was significantly upregulated. These data show that CM and MM models have a distinct gene expression signature including an AP-2 transcription factor switch and decreased MAPK activation as shown by pERK staining and MAPK target gene expression.

**Figure 2:**
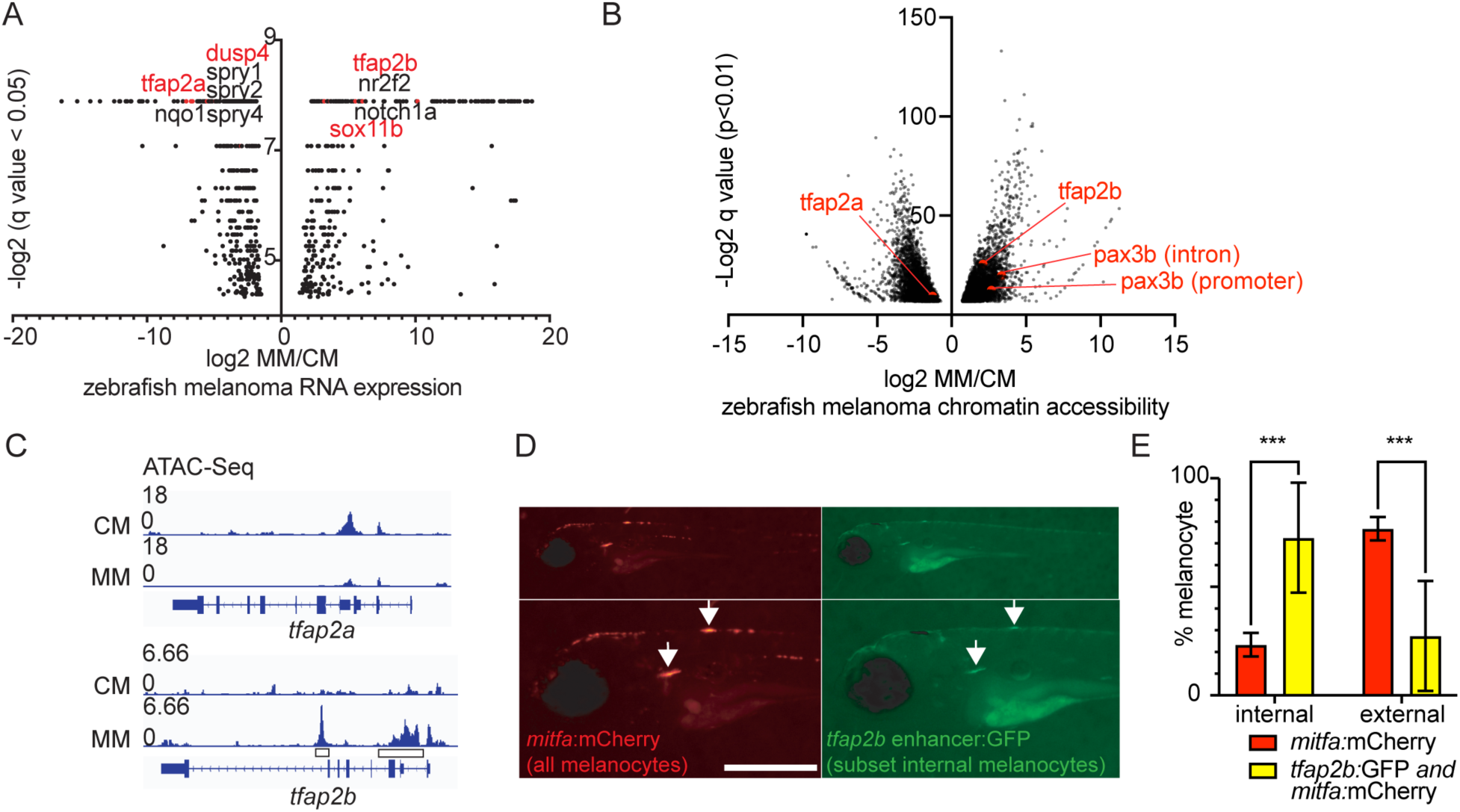
Zebrafish MM has a distinct cellular state from CM. A-B) Volcano plots of A) bulk RNA-seq and B) ATAC-seq from zebrafish MM vs. CM. C) IGV plot of ATAC-seq showing chromatin accessibility at *tfap2a* and *tfap2b* in zebrafish MM vs. CM. Boxes = loci used for *tfap2b* reporter. D-E) *tfap2b* enhancer-driving GFP preferentially labels internal *mitfa*-mCherry labeled melanocytes in 6-day old Casper zebrafish. D) Fluorescent images. Scale bar = 500um. E) Reporter expression quantification. p-value = 0.0003, 2-way ANOVA with multiple comparisons.

To determine whether the internal melanomas had a distinct epigenetic state, we performed the assay for transposase-accessible chromatin using sequencing (ATAC-seq) on MM (n=3) vs. CM (n=3). MM had 4901 significantly more accessible peaks and 4719 less accessible peaks compared to CM (q<0.01) (Fig. 2B). As we previously observed a switch in AP-2 transcription factor gene expression, we asked whether there was also a change in chromatin accessibility at the loci near these genes. The chromatin surrounding *tfap2a* was open in CMs but closed in the MMs, while the chromatin surrounding the related transcription factor *tfap2b* was closed in CMs but open in MMs (Fig. 2C). *tfap2b* was recently shown to be a marker of melanocyte stem cells in zebrafish^18^, suggesting that MM may derive from a less-differentiated melanocyte population. We also found that the early neural crest transcription factor *pax3b,* which has been shown to maintain melanocytes in mice in an undifferentiated state^19^, had proximal loci that were significantly more accessible in MM than in CM (Fig. S2B).

As the *tfap2b*-proximal loci were consistently open in MM model and closed in CM, we wondered if we could use the *tfap2b*-proximal sequences to drive GFP to label internal melanocytes. The two most statistically enriched *tfap2b*-proximal sequences (boxes in Fig. 2C) that were present in all MM samples and absent from all CM samples were cloned preceding a *beta-globin* minimal promoter and GFP. To test whether *tfap2b-*proximal sequences could label internal melanocytes, we expressed the *tfap2b* reporter in zebrafish embryos. The *tfap2b* reporter was injected into Casper zebrafish with a vector that labels all melanocytes red (*mitfa-*mCherry) and reagents to CRISPR-delete *tyrosinase* to remove pigment to allow fluorescent imaging (schematic Fig. S2C). Injected fish were imaged at 6 days post fertilization (dpf) and the *tfap2b-* GFP reporter labeled a subset of internal and external melanocytes (Fig. 2D). Significantly more double labeled (*tfap2b-*GFP*; mitfa-*mCherry) melanocytes were identified inside the zebrafish (Fig. 2E). These data show that zebrafish MM has a unique epigenetic state and that a MM enhancer can be used to label a subset of normal internal melanocytes that are amenable to developing MM.

### Zebrafish internal melanocytes have properties consistent with MM initiating cells

To identify the location of the melanocytes that could be initiating MM in zebrafish, Casper zebrafish with melanocyte-specific GFP expression were generated and grown to adulthood. The abdominal organs were removed to visualize melanocytes in the abdominal cavity. Brightfield and fluorescent microscopy demonstrated that there were three different morphologic populations of pigmented cells on the inside of the zebrafish including those with a dendritic morphology in the anterior abdomen near the head, many that lined the superior aspect of the kidney marrow as previously reported^20^, and those lining the abdominal cavity (Fig. 3A). Melanocyte-containing zebrafish skin or internal tissues were dissected, mechanically dissociated, digested into a single cell suspension, and fluorescence activated cell sorted (FACS) for GFP. The internal or cutaneous melanocytes were sorted and subjected to single-cell RNA sequencing (scRNA-seq) using the SORT-seq platform^21^, which allows matching of gene expression with FACS data and enhanced transcript detection (Fig. 3B). Sorted cells from *mitfa:*GFP zebrafish had GFP expression vs. negative controls (Fig. S3A-B). Gene expression was normalized, and cells were clustered using Seurat^22^ (Fig. 3C and Fig. S3C). Cells expressing *mitfa,* the master transcriptional regulator of the melanocyte lineage, were hypothesized to represent true melanocytes, and informatically isolated (Fig. 3D) and re-clustered (Fig. 3E). Three distinct melanocyte populations were observed. One melanocyte population derived from only inside the zebrafish (labeled “internal”). Two other groups of melanocytes were observed, one almost entirely from zebrafish skin (labeled “external”) and another that contained cells from both locations (labeled “both”). We asked whether any of the melanocytes in Fig. 2E expressed *tfap2b* vs. *tfap2a.* The “external” and “both” melanocyte groups expressed *tfap2a* (Fig 3F, first panel); whereas a subset of the “internal” melanocytes expressed *tfap2b,* consistent with the *tfap2b* reporter identifying a subset of internal melanocytes as in Fig. 2C-D (Fig. 3F second panel). To determine which melanocyte population likely was the cell of origin for MM, we asked which normal melanocyte population had the most similar gene expression to zebrafish MM tumors. “Internal” melanocytes alone expressed all 12 genes that were upregulated in zebrafish MM from Fig. 2A, many at high levels and in a large percentage of cells (Fig. 3G, left). “Internal” melanocytes also expressed high levels of cell cycles genes including those that are typically expressed in actively dividing cells (S or G2/M phase), indicating that these cells may represent a transit amplifying melanocyte population in adult zebrafish (Fig. 3G, see Cell Cycle Genes, Fig. S3D). These data show that internal adult zebrafish mucosal melanocytes have a distinct cell state and that internal melanocytes have gene expression profiles consistent with being the cell of origin for zebrafish MM.

**Figure 3:**
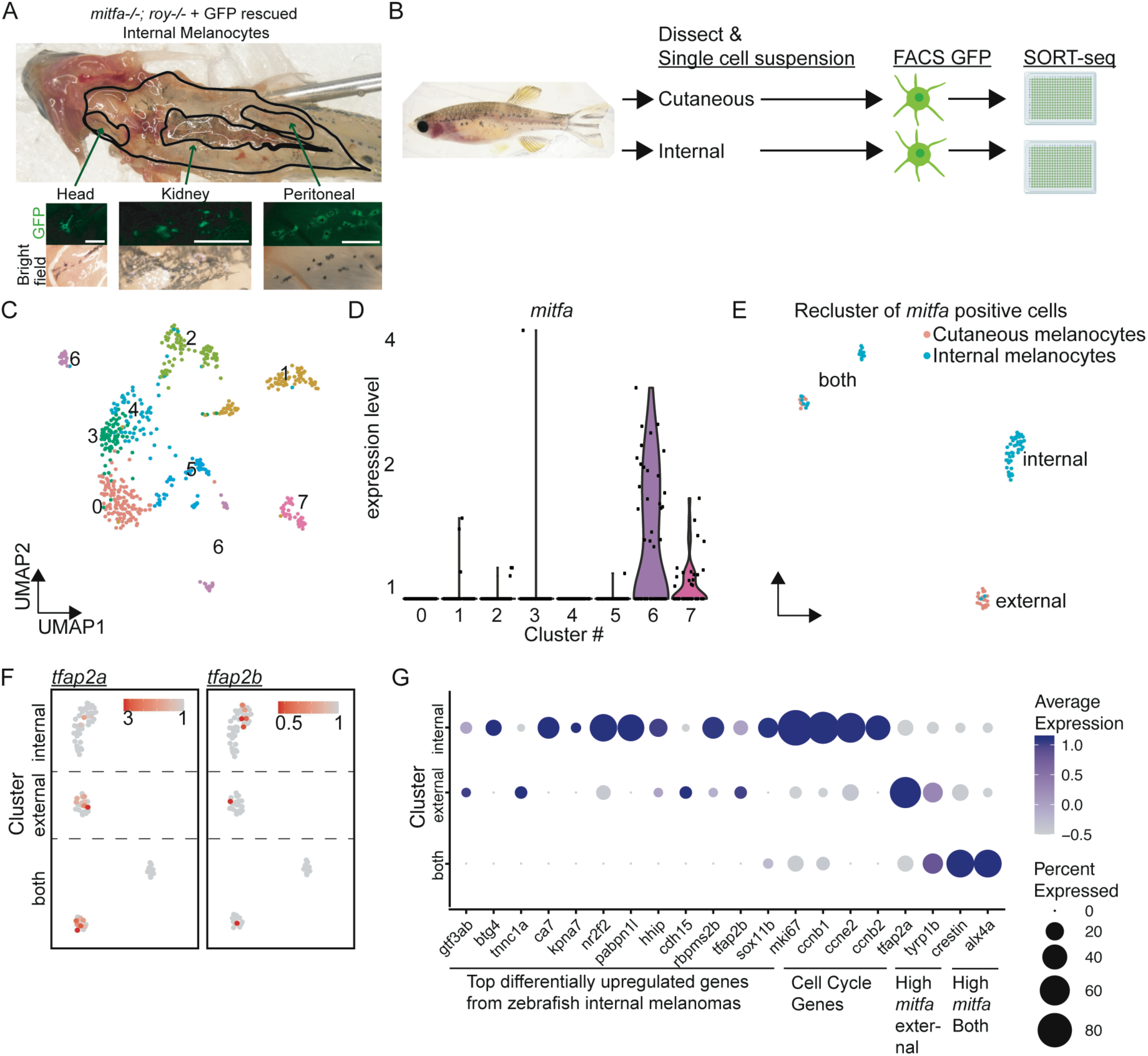
Zebrafish internal melanocytes have properties consistent with MM initiating cells. A) Light micrographs demonstrating internal GFP-expressing melanocytes from adult Casper zebrafish. Scale bar = 500um. B) Schematic for single-cell RNA-seq (scRNA-seq) of cutaneous and internal adult zebrafish melanocytes. C) UMAP-plot of scRNA-seq from GFP-sorted external and internal melanocytes. D) *mitfa-*expressing cells from C). E) UMAP of re-clustered *mitfa*+ melanocytes. Blue = internal. Red = external. Labels = cluster names. F) *tfap2a* and *tfap2b* expression in “internal”, “external”, and “both” melanocytes. G) Dot plot showing gene expression of top differentially expressed genes from zebrafish MMs for “internal”, “external”, and “both” melanocytes from E).

### MM cell state is conserved in patients

To determine whether the zebrafish model accurately reflects the expression programs and heterogeneity of human MM, we completed single-cell RNA-seq (scRNA-seq) of patient MM (n=10) and compared to published scRNA-seq from patient CM (n=23) ^23^ (Fig. 4A, S4A, clinical data in Data S2:Tab 1-2). Patients had a similar average age (median age 68yrs, MM and CM) (Fig. S4B). A higher proportion of MM patients were female (60% MM vs. 35% CM) (Fig. S4C) and a similar percentage were from metastatic vs. primary lesions (87% CM vs. 80% MM) (Fig. S4D). Of the MM patients, four were sinonasal, four anal, and two vulvovaginal. Targeting sequencing panels from two different institutions were consistent with published MM genomics reports^3, 4, 5, 6^. For example, two of four nasal MM patients had activating NRAS mutations (Q61K and G12A)^6^, two of the anal MM patients harbored SF3B1 hotspot mutations (R625C and R625C)^24^, and two patients had activating KIT mutations and gain of the KIT locus (N822K and L576P)^25^ (Fig. S4E, Data S2:Tab 3). As our zebrafish MM model was built using tumor-cell specific genetic changes, we focused on tumor cell intrinsic cell states in patient MM by informatically isolating tumor cells. Consistent with MM having distinct chromatin state from CM, a UMAP-plot showed that MMs clustered separately from CMs (Fig. 4A). We also observed that MMs were spread more broadly on this UMAP than CMs, suggesting that gene expression in MM was more heterogenous than in CM as expected. These data show gene expression characteristics of MM are distinct from CM.

**Figure 4:**
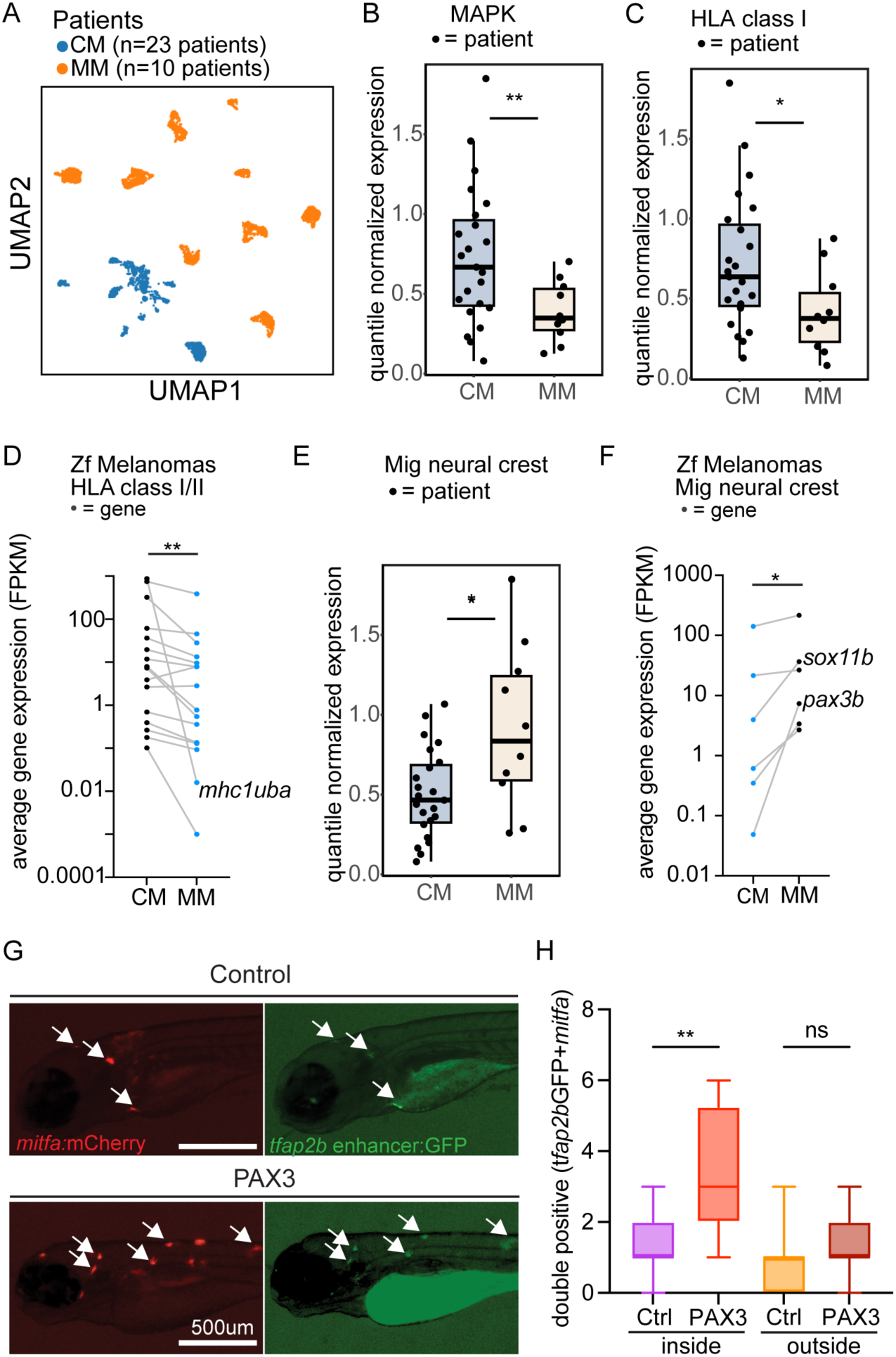
MM cell state is conserved in patients. A) UMAP-plot of single-cell RNA-seq (scRNA-seq) from patient tumors, CM patients (n=23) and MM patients (n=10). B-C) Mean normalized gene expression in CM vs. MM cells from scRNA-seq for B) MAPK target genes, p-value = 0.0036 two-sided t-test and C) HLA Class I antigen presentation genes. p-value = 0.017 two-sided t-test. Dots = patients. D) Average gene expression of expressed zebrafish *mhc class I/II* genes in a zebrafish CM vs. MM. Dots = genes. Lines indicate the same gene in two conditions. p-value = 0.0017, two-tailed Wilcoxon matched-pairs rank test. E) Mean normalized gene expression in MM vs. CM cells from scRNA-seq for migratory neural crest genes. p-value = 0.036 two-sided t-test. F) Average gene expression (FPKM) of migratory neural crest genes in zebrafish CMs compared to the MMs. Dots = genes. Lines indicate the same gene in two conditions. p-value= 0.031, two-tailed Wilcoxon matched-pairs rank test. G-H) Melanocyte-specific expression of hPAX3 vs. control in zebrafish with labeled internal melanocytes (double positive *mitfa-mCherry* and *tfap2b-GFP)* and pigment removal at 5dpf. G) Representative immunofluorescent images. Scale bar = 500μm. H) Quantification of double positive melanocytes inside and outside zebrafish embryos. p-value = 0.0014, ordinary one-way ANOVA with multiple comparisons.

As the zebrafish MMs had downregulation of MAPK gene expression, we hypothesized that MAPK target gene expression would also be lower in patient MMs vs. CMs. Expressed MAPK target genes ^26^ (*SPRY2*, *SPRY4*, *ETV4*, *ETV5*, *DUSP4*, and *DUSP6*) were assessed in MM vs. CM. As expected, MAPK target gene expression was significantly lower in MM (p = 0.0036, two-sided t test) (Fig. 4B and Fig.S4F). Lower MAPK target gene expression is concordant with MM patients harboring fewer and less potent MAPK activating mutations as published^6, 27^. We also observed fewer and less potent MAPK activating mutations in our data (scRNA-seq patients, 0/10 had BRAF mutations and 2/10 had NRAS mutations) (Data S2:Tab 3). In our larger cohort, 8% (7/86) had BRAF mutations with 71% of those being non-V600 (Fig. S4G-H, Table S3, Data S1); while 16% had NRAS mutation (n=14/86, Data S1). These data show that MM patients, like MM zebrafish, have lower MAPK pathway target gene expression. This illustrates that MM exists in a cellular state that is less reliant on MAPK pathway activation.

To determine the unbiased gene expression changes in MM vs. CM, differential expression analysis of pseudobulked scRNA-seq of MM vs. CM was completed. This analysis identified that the most downregulated pathway was “Interferon Signaling” (Fig S4I-J), and this gene set included multiple HLA class I genes. The average pseudobulked expression for four HLA class I genes (HLA-A, HLA-B, HLA-C, B2M) was compared between MM and CM and identified that HLA class I gene expression were significantly downregulated in patient MM (p = 0.017, two-sided t-test) (Fig. 4C and S4K). We wondered if the decreased expression of MHC class I genes was conserved between patients and the zebrafish model. All expressed zebrafish *mhc* genes were plotted (FPKM >= 0.1 in at least one condition) for the zebrafish MM and CM models. Collectively *mhc* gene was expressed at lower levels in the zebrafish MM model as compared to the CM model (p=0.007, Wilcoxon 2-sided t-test) (Fig. 4D). We hypothesize that the decreased expression of *mhc* genes is occurring in MM tumor cells because we saw no change in expression in genes that uniquely identify T-cell genes in zebrafish (Fig. S4L). These results suggest that the MM cell of origin has a cell state with low MHC class I expression, and that this cell state is maintained in melanomas derived from mucosal melanocytes. Lower MHC class I expression in MM could contribute to the lower observed immune therapy responses observed in MM patients.

To understand what drives the MM cell state, we asked what pathways define MM. The zebrafish MM model revealed differential expression of genes involved in normal melanocyte differentiation from their developmental neural crest precursor cells. We asked whether there was differential expression in MM vs. CM of genes that define premigratory vs. migratory cranial neural crest cells (CNCC)^28^. Migratory CNCC genes were significantly upregulated in MM vs. CM (*PAX3*, *SOX9*, *SOX10*, *TFAP2A*) (p = 0.036, two-sided t-test) (Fig. 4E), while premigratory CNCC genes were not differentially expressed (*SNAI1*, *SNAI2*, *MYC*, *MYCN*, *ETS1*) (p = 0.97, two-sided t test). Zebrafish MM RNA-seq also showed increased zebrafish migratory neural crest gene expression (Fig. 4F). The specific expression of *tfap2b* or *sox11b* that was observed in zebrafish MM, was not conserved in patients as both TFAP2B and SOX11 were rarely expressed in MM or CM. As we observed that *PAX3* was upregulated in patient MM and in zebrafish MM; and as the ortholog *pax3b* had loci with increased chromatin accessibility from zebrafish MM ATAC-seq, we investigated whether PAX3 could play a conserved role in contributing to the cell state of mucosal melanocytes.

PAX3 is a gene required for melanocyte development in humans as loss of PAX3 is associated with a genetic condition that causes hearing loss and changes in the pigmentation of hair, eyes, and skin^29^. Pax3 expression in mice has been shown to maintain melanocytes in an undifferentiated state^19^ and PAX3 expression in human melanomas is associated with fewer immune therapy responses and worse overall survival of patients in two cohorts ^30^. We asked whether human PAX3 (hPAX3) expression in zebrafish could enrich for internal melanocytes as measured by our internal melanocyte reporter built in Fig. 2. hPAX3 or a control vector were expressed in a melanocyte-specific manner in Casper zebrafish in addition to the reporter constructs. Internal melanocytes (double positive for *tfap2b:GFP* and *mitfa:*mCherry) and external melanocytes (single positive for *mitfa:*mCherry) were imaged as in Fig. 2. hPAX3 expression caused significantly more internal melanocytes (Fig. 4G-H) without affecting the total number of melanocytes observed (Fig S4M), indicating that PAX3 expression is one factor that can functionally enrich for mucosal melanocytes. As our data suggests that PAX3 is functionally contributing to the MM cell state, we hypothesized that open chromatin from MM would be enriched for PAX3 binding sites. The binding sequence for PAX3 was significantly enriched in ATAC-seq peaks that are more accessible in zebrafish MM than in CM (E value=8.45e-15, SEA). These data suggest that PAX3 is a conserved contributor to the cell state of MM, which has unique oncogene susceptibility and that has features predicting for its lethal clinical behavior.

## Discussion

We developed a novel MM model in zebrafish that faithfully recapitulates both the transcriptional and histological features of human MM. To our knowledge, this is the first genetically engineered model of this disease. Using this model, we identify a distinct melanocyte population inside zebrafish that is uniquely susceptible to malignant transformation upon introduction of oncogenes and loss of tumor suppressors as found in MM patients. The cell of origin for MM appears to be a developmentally distinct melanocyte population that relies less on MAPK signaling and has muted antigen presentation programs relative to cutaneous melanocytes. We find that both the zebrafish MM model and MM patients have increased migratory neural crest gene expression, which could explain the more metastatic nature of these tumors in patients. We also find that the zebrafish model and patients have decreased tumor-cell-intrinsic expression of antigen presentation genes, which could explain why these patients have a lower response rate and fewer durable responses with immune therapy. Our comparative analyses reveal striking conservation between the genetics and cell states underlying zebrafish and human MM. These findings establish this zebrafish model as a representative system to uncover drivers and drug targets for this rare and lethal disease in future work.

We have generated the first genetically engineered model of MM. While there are current efforts to establish MM human cell lines^31, 32^ and PDX models and it was previously known that dogs naturally are at risk for MM^33, 34, 35^; genetically engineered models can offer important information about causality and are more experimentally tractable. For instance, we observed decreased tumor-cell-intrinsic MHC class I gene expression in patient MMs, which could be caused by the cell state of the tumor cells vs. secondary effects from the tumor microenvironment. Our zebrafish model offers clarity. As we induced MM in zebrafish using melanocyte-specific genetic changes and observed MMs with decreased MHC class I expression, our data suggest that cell autonomous genetic changes in mucosal melanocytes drive a cell state that has decreased MHC class I expression.

In MM patients, we observed a significant downregulation of MAPK target and antigen presentation gene expression as compared to CM. Our observation that MMs have lower MAPK signaling aligns with the lack of frequent strong activating MAPK mutations found in MM patients, suggesting less reliance on the MAPK pathway for transformation relative to CM. Lower MHC class I expression in our study also fits with poorer immunotherapy responses seen clinically in MM patients. We instead find MMs have a conserved cell state that expresses migratory neural crest genes. In fish, we were able to label a subset of internal melanocytes using a reporter built from MM-specific *tfap2b* enhancer sequences. Our reporter is distinct from a prior *tfap2b* reporter that includes the promoter sequences and labels melanocyte stem cells^18^. As TFAP2B expression was not conserved in MM patients, we chose instead to test for a functional role of PAX3 in MM. Our results suggest that MM arises from a different melanocyte pool that is localized inside animals.

PAX3 is a transcription factor that plays a role in neural development and is required for melanocyte differentiation. Mutations in PAX3 in humans cause Waardenburg syndrome type 1 which is associated with pigmentation defects^36, 37, 38, 39^. In mice, PAX3 has been shown to act at a nodal point in melanocyte stem cell differentiation, where it is required for the melanocyte stem cell state while also priming these cells for differentiation^19, 37^. We found that PAX3 expression in zebrafish can contribute to increased numbers of internal melanocytes, suggesting a functional role for PAX3 in specifying the cellular state of internal melanocytes. As PAX3 is part of the migratory neural crest program, perhaps MMs derive their innate capacity for metastasis from continued expression of the migratory neural crest program. While PAX3 is not amplified in melanomas, its sustained expression is associated with worse survival rates in patients with melanoma^30, 40^. Our data and the literature nominate PAX3 as one potential functional mediator of the metastatic-like phenotype of MM.

Cell state can predict for oncogene susceptibility. Our data suggest that the MM cell of origin is an internal melanocyte that is uniquely susceptible to genetic changes that occur in MM, i.e. expression of CCND1 and loss of *PTEN* and *TP53*. Although all melanocytes in the zebrafish model have the aforementioned genetic changes, only the internal melanocytes are transformed into melanoma; while the cutaneous melanocytes are spared. Our work complements the work of Weiss et. al.^13^ that found that zebrafish fin melanocytes, analogous to human acral melanocytes, are enriched for transformation by an oncogene amplified in acral melanoma. Together our studies illuminate that there are distinct melanocyte cell states that relate to their anatomic location and predict their sensitivity to different oncogenic drivers. The cell state of the originating melanocyte ultimately influences the clinical behavior of derived melanomas, including treatment responses.

Our work shows that by modeling genetic changes in MM patients, genetically engineered MM models can be built. Our work suggests that the cell of origin for MM is distinct from the cell of origin for CM. MMs exist in a state that is less reliant on MAPK signaling (and thus less responsive to MAPK pathways inhibitors) and have less MHC class I expression, which could contribute to the fewer and less durable immune therapy responses observed for MM. In future work, it will be important to discover agents that are either lethal to this cell state or can rescue MHC class I expression, with the hope of rescuing durable immune therapy responses in these patients. We hope that this model and subsequent iterations will help to define the functional genetic changes in MM to inform the development of mouse models, drive therapeutic discovery, and ultimately focus future clinical trials for MM patients.

## Supplementary Materials

Materials and Methods

Figs. S1 to S4

Tables S1 to S3

References 1-36

Data S1 to S2

## Acknowledgments

The authors would like to acknowledge the DFCI Oncology Data Retrieval System (OncDRS) for the aggregation, management, and delivery of genomic research data used in this project.

## Data and materials availability

The datasets generated and/or analyzed during the present study have been uploaded to the Gene Expression Omnibus (GEO pending). All zebrafish strains, and DNA vectors are readily available through the corresponding author. Where possible, we can share remaining zebrafish tumor material; however, this material is limited in abundance. All antibodies are commercially available.

