## Supplemental Materials for "Specific oncogene activation of the cell of origin in mucosal melanoma"

**The PDF file includes:**

Materials and Methods
Figs. S1 to S4
Tables S1 to S3
References 1 to 36

**Other Supplementary Materials for this manuscript include the following:**

Data S1 to S2

**Materials and Methods**

**Zebrafish:**

*Cloning:* For the *tfap2b-GFP* reporter, two open areas of chromatin that were statistically enriched in mucosal melanoma (MM) as compared to cutaneous melanoma (CM) from ATAC-seq (see ATAC-seq below) were concatenated and TOPO cloned into a 5' entry vector (Invitrogen, K59120). The 5' entry vector was recombined using LR clonase (Invitrogen, 12538-120) with a mouse Beta-globin minimal promoter[1] fused to eGFP and a polyA tail, into a MiniCoopR (MCR) vector flanked by Tol2 sites. Human PAX3 (hPAX3) coding sequence was amplified from Addgene plasmid 176360. hPAX3 sequence was recombined with EcoR1/Sal1 cut MCR[2] in a Gibson reaction (NEB E2611).

*Melanoma models:* Animal studies were approved by Dana Farber Animal Care and Use Committee (Protocol IACUC 21-027) or by Boston Children's Hospital (Protocol IACUC 20-10-4253R). Experiments were performed as published[3, 4, 5]. Briefly, *roy*$^{-/-}$;*mitfa*$^{-/-}$ zebrafish[6] (hereafter Casper) one-cell embryos were microinjected with 20 ng/uL DNA along with tol2 *in vitro* transcribed RNA for integration. CM was modeled by injecting equal amounts of CRISPR MCR:*tp53 gRNA*[7] and MCR:hBRAF$^{V600E}$[7], whereas MM was modeled by injecting equal amounts of MCR:hCCND1[8], CRISPR MCR:*pten a/b gRNA*[7] and CRISPR MCR:*tp53 gRNA*[7]. Embryos were sorted for melanocyte rescue at 5 days post fertilization (dpf) to ensure vector integration. 20 zebrafish were raised per tank to control for density effects. Zebrafish were scored visually for the development of internal (internal mass protruding >1mm$^2$) and external melanomas (external mass >1mm$^2$). Melanoma-free survival curves and Log-rank tests were generated in GraphPad Prism. Representative adult zebrafish with CM and MM were anesthetized in tricaine and imaged using brightfield imaging.

*Tfap2b reporter*: To visualize internal melanocytes, *tfap2b:GFP* was injected into Caspers as above and assessed in the first generation. To label melanocytes red, *mitfa:mCherry* was co-injected. To improve imaging, pigment was removed via CRISPR-deletion of *tyrosinase (tyr)* (10ng of *in vitro* transcribed *tyr* gRNA + 200ng of Cas9 protein) (PAN Bio, CP01) (Schematic in Fig. S2C). For Fig. 2, 6dpf embryos with expression of both transgenes and disrupted eye pigmentation (demonstrating functioning *tyr* CRISPR) were selected and fluorescently imaged on a Nikon SMZ18. For each fish, the location (i.e. internal or external) of melanocytes (*mitfa*:mCherry positive*)* that either had or did not have *tfap2b:GFP* expression was recorded. For hPAX3-expression experiments in Fig. 4, the two reporters were injected along with MCR:*mitfa:*PAX3 or a control MCR:*mitfa:*empty vector. Injected embryos were incubated at 28°C for two days and then incubated at room temp for 3 days. On day 6 they were sorted for presence of both reporters and functional *tyr* CRISPR-deletion as above and fluorescently imaged. The location of *mitfa* positive (red) and double positive melanocyte (red and green) was recorded.

*Immuno Histo Chemistry:* Zebrafish with MM and CM were fixed in 4% paraformaldehyde and paraffin embedded, sectioned, stained with primary antibodies (anti-CCND1, ab134175; anti-pERK, CST 4370S), and then stained with secondary antibody for brown IHC (Leica, DS9800). Zebrafish tumors were scored for presence/absence of CCND1 and pERK.

*CRISPR cutting verification:* Melanoma CRISPR-deletion (using gRNAs in Table S1) was verified via DNA extraction, PCR across the cut site (primers used in Table S2), and NGS of PCR reads (sequenced using: https://dnacore.mgh.harvard.edu/new-cgi-bin/site/pages/crispr_sequencing_main.jsp). Data sets were mapped to the *Danio rerio* genome (version GRCz11) using Bowtie (version 0.12.9). CrispRVariants (version 3.18)[9] was used to identify insertions and deletions (indels) around the gRNA site. Indels at the gRNA cut site in >2% of reads were used in downstream calculations. Reads that were predicted to maintain function included in-frame indels as well as wild type reads, while out-of-frame reads were predicted to cause loss of function.

*RNA-Seq:* MM (Casper fish injected with MCR:CCND1; CRISPR MCR:*pten a/b sgRNA*; CRISPR MCR:*tp53 sgRNA*) or CM (*mitfa*-/-; *tp53*-/-; BRAF$^{V600E}$ fish injected with MCR:GFP) were collected. Tissue was disrupted using QIAshredder columns (Qiagen, 79656). DNA was removed and RNA was purified using columns (Qiagen, 74134). polyA-selected RNA (NEB, E7490S) RNA libraries were prepped with random priming (NEBNext Ultra RNA Library Prep Kit for Illumina, E7530) and paired ends were sequenced on a Novoseq 6000.

*RNA-Seq analysis:* Quality control of RNA-Seq datasets was performed by FastQC (https://www.bioinformatics.babraham.ac.uk/projects/fastqc/) and Cutadapt[10] was used to remove adaptor sequences and low quality regions. High-quality reads were aligned to UCSC build danRer11 of zebrafish genome using Tophat 2.0.11[11] without novel splicing form calls. Transcript abundance and differential expression were calculated with Cufflinks 2.2.1[12]. FPKM values were used to normalize and quantify each gene. Significantly changed genes ($q<0.05$) were plotted in Fig. 2A. Average FPKM was plotted for relevant genes in CM vs. MM for Fig. 4D and 4F.

*ATAC-Seq:* Tumors were harvested and prepared for transposition reaction following the manufacturers protocol (Nextera 15028212). 5,000 sorted melanocytes (*mitfa*:mCherry positive*)* from 3 combined CM (*mitfa*-/-; *tp53*-/-; BRAF$^{V600E}$; *crestin*:GFP fish injected with *MCR:empty*, *mitfa:*mCherry, *tyrosinase gRNA* and Cas9) or 40,000 cells from each MM (Casper fish injected with MCR:CCND1; CRISPR MCR:*ptena/b sgRNA*; CRISPR MCR:*tp53 sgRNA*) were lysed and subjected to tagmentation reaction and library construction as previously described[13]. Libraries were run on an Illumina HiSeq 2500 or a NovaSeq 6000.

*ATAC-Seq analysis:* To identify a uniform set of regions-of-interest, ATAC-Seq reads were aligned to the primary chromosomes of the danRer11 genome build, i.e. chr1-chr25, using bowtie v1.2.2[14] in single-end mode with parameters -k 1 -m 1 –best and -l set to the read length. Mapped reads from technical replicate samples were combined. Enriched regions were called using MACS v1.4.1[15] with -p 1e-9. To create a unified set of regions-of-interest, peaks were collapsed across all six biological samples using bedtools merge[16]. Collapsed peaks were assigned to the RefSeq gene (downloaded 8/6/20) with the single most proximal transcription start site using bedtools closest. To quantify coverage genome-wide and in regions of interest, ATAC-Seq reads were trimmed using cutadapt[17] with parameters –minimum-length 25 -a CTGTCTCTTATACACATCT -A CTGTCTCTTATACACATCT. Trimmed reads were aligned to the primary chromosomes of the danRer11 genome build, i.e. chr1-chr25, using bowtie v1.2.2 in paired-end mode with parameters --allow-contain --maxins=100000 --chunkmbs=256 -k 1 -m 1 –best. Fragments were generated from paired-end mappings using samtools[18] view, samtools sort, bedtools bamtobed, and bedtools bedtobam. These fragments were used to calculate coverage of collapsed peaks defined above using bedtools intersect -c. Fragment counts in each collapsed region were used as input for

DEseq2[19] with default parameters to compare the cutaneous and mucosal region coverage. Genome-wide coverage in 50bp bins was quantified using bedtools intersect -c, normalized to the number of mapped reads per sample, and converted to TDF with igvtools[20]. Significantly changed chromatin accessibility peaks were plotted ($q<0.01$). The DEseq2 output for ATAC-seq differential enrichment between mucosal and cutaneous melanomas was used to define sets of MM-specific and CM-specific peaks with adjusted p-values < 0.05 and log2 fold-changes of > or < 0. 100bp regions centered on the midpoints of these peaks were used for motif enrichment analysis. The sequences of these 100bp regions in the danRer11 genome were acquired using bedtools getfasta and used as input for SEA (https://www.biorxiv.org/content/10.1101/2021.08.23.457422v1) with the CIS-BP_2.00 Danio rerio position-weight matrices and using the sequences from CM-specific peaks as a control.

*SORT-seq:* To image internal melanocytes, *mitfa*:GFP expressing Caspers were grown until 13.1 weeks old and sacrificed. To image internal melanocytes, the peritoneal cavity was opened, all internal organs were removed except for the kidney marrow because melanocytes line this organ. Brightfield and GFP images were captured. The internal tissues with melanocytes were dissected from seven fish and combined. For external melanocytes, two fish were skinned and the skins were combined. All tissues were mechanically dissociated, digested in TrypLE for 45 min (Invitrogen, 12563011), 40uM filtered, spun down, and resuspended in FACs buffer (dPBS Mg+/Ca+ free, with 2% FBS, pen/strep). Sytox blue was added at 1:1000 to label dead cells. Live GFP-positive cells were FACs sorted using a BD FACSAria. 192 GFP-positive cells from zebrafish skins were sorted into one plate and 158 GFP-positive cells from zebrafish internal tissues were sorted into another plate. At least 8 wells per plate were retained with no cells as negative controls. Plates were provided by Single-Cell Discovery (https://www.scdiscoveries.com/), and each well was pre-filled with 10ul mineral oil containing a 50nl droplet of uniquely barcoded primer for polyA capture of mRNA. Plates with sorted cells were spun at 1000g for 1 minute to ensure that droplets containing the single cell and barcoded primers would merge into a single drop. Plates were stored at -80C and shipped on dry ice to Single-Cell Discovery for library preparation using an adapted version of the SORT-seq protocol[21]. Briefly, cells were heat-lysed at 65°C followed by cDNA synthesis. Next, all the barcoded material from one plate was pooled into one library and amplified using in vitro transcription. Following amplification, library preparation was done following the

CEL-Seq2 protocol[22] to prepare a cDNA library for sequencing using TruSeq small RNA primers (Illumina). The DNA library was paired-end sequenced on an Illumina Nextseq™ 500, high output, with a 1×75 bp Illumina kit (read 1: 26 cycles, index read: 6 cycles, read 2: 60 cycles).

*SORT-seq analysis:* BWA was used to align paired-end reads to the zebrafish genome danRer11[23] and data was demultiplexed as described in[24]. Demultiplexed datasets were mapped, and count tables were generated using MapAndGo. Count tables were corrected using UMI to remove duplicate reads. Transcript counts were adjusted using Poissonian counting statistics to yield the number of UMIs detected per cell. Counts were imported into R using the Seurat suite version 3.0[25]. Data was filtered using the following parameters: >100 transcripts per cell, present in at least two cells, <10,000 transcripts, <50% mitochondrial transcripts, and 20 principal components.

**Patient data**

*Genetic analysis:* Mutation and copy number alteration data for CDK4 and CCND1 from CM patients with available GISTIC and mutational analysis (n=363)[26] was downloaded from cBioPortal (www.cbioportal.org) on 11/16/23 and this analysis was described in the text.

*Targeted sequencing and co-mutation analysis:* Tumor genotyping of DFCI MM patients (n = 86) was performed by targeted sequencing of up to 447 genes (Oncopanel) of 59 MM patients that were previously published[27] in addition to 27 newly sequenced patients. The following variant types were assessed and included if present: LA (low amplification), HA (high amplification), 1DEL (loss of one copy), 2DEL (loss of both copies), Missense, Splice, Structural Variation. Indel, Nonsense, and Truncation were grouped and labeled as "Others". Python package CoMut[28] was used to plot the mutations. Samples with three or more variants on one gene are indicated as "Multiple". Only samples with at least one alteration present were displayed. Lollipop plot of BRAF mutations was generated using R package Maftools. Customized function adapted from function lollipopPlot(). For DFCI single-cell RNA-seq patients, data is shown for 22 genes that were previously shown to be mutated in MM. For MGH patients single-cell RNA-seq patients, all mutations detected by Snapshot targeted sequencing were included.

*Single-cell RNA-seq (scRNA-seq) sample processing:* IRB approval was obtained prior to study enrollment and written informed consent was obtained from the patient for the collection of tissue and blood samples for research and genomic profiling, as approved by the Dana-Farber/Harvard Cancer Center Institutional Review Board (DF/HCC Protocol 11-181 for MGH patients or 05-042 for DFCI patients). All MM samples were disassociated, frozen, then thawed for 10x scRNA-seq.

*scRNA-seq analysis:* For scRNA-seq samples in CM, we downloaded the raw count matrix from Jerby Arnon[29] (GSE115978) and subset for tumor cells based on the original cell type annotations from this study. Patient samples without tumor cells were excluded from analysis (8 out of 31). For MM samples, we used Cell Ranger (v3.1) to perform sample demultiplexing, barcode processing, alignment (reference genome hg19 to match Jerby Arnon dataset), filtering, and UMI counting. Cellbender[30] was used for ambient RNA correction. Python package Scrublet[31] was used for doublet removal. Python package scanpy (v1.9.1)[32] was used for subsequent data processing and analysis. Briefly, cells with fewer than 200 genes or more than 6,000 genes, and those with greater than 25% mitochondrial genes were removed from each sample. 4,000 most highly variable genes from the normalized counts were used for principal components analysis (PCA). Following PCA, Uniform Manifold Approximation and Projection (UMAP) was performed on the first 25 principal components and 100 nearest neighbors to visualize preliminary cell clusters. Unsupervised clustering using the default graph-based UMAP algorithm implemented in scanpy (resolution parameter 0.3) identified 13 distinct clusters. Based on differentially expressed genes among clusters, tumor cells (S100A1, S100B, MITF, PMEL, MLANA, SOX10) were separated from other cell types including immune cells (PTPRC, CD3E, LYZ) and endothelial cells (CLEC14A, CD34). Non-tumor cells were removed. Remaining tumor cells with higher than 15% mitochondrial genes were removed to further clean up the data. We then merged malignant MM tumor cells 10X data with CM tumor cells smartseq2 data, yielding a matrix containing data from 38 samples (23 CM, 15 MM) from 33 patients (23 CM, 10 MM). To mitigate technical variation between 10X and Smart-seq sequencing platforms, we included only genes detected from both datasets for analysis, and downsampled smart-seq samples to 5,500 counts per cell (approximately the median of the counts for 10X MM samples). Only genes expressed in at least 0.5% of cells were kept for analysis. We normalized the data, selected top 4,000 most highly variable genes for PCA, and used the first 25 principal components and 50 nearest neighbors for UMAP visualization.

For UMAP visualization purposes, downsampling was performed to include at most 500 cells per patient. For analysis, all cells were included. MAPK transcriptional targets were as published[33] (SPRY2, SPRY4, ETV4, ETV5, DUSP4, and DUSP6). CCND1 was excluded as it can be amplified independent of MAPK induction (as shown in Fig. 2A) and EPHA2 and EPHA4 were excluded due to their expression in a small proportion of cells (<10%). "mig_cncc" and "premig_nc" gene sets were as published in[34]. We scored single cells with the above custom gene signatures based on pathway biology ("mapk", "classI_AP", "mig_cncc", "premig_nc") using scanpy function sc.tl.score_genes(). We then generated pseudobulk data by calculating the mean of genes and pathway scores per cell by patients. To mitigate the potential biases from platform-specific efficiencies in sequencing RNA, which is particularly shown for genes expressed at extreme levels which can be seen with mitochondrial genes in 10x vs. Smart-seq[35], we performed quantile normalization on the pseudo-bulk RNA data. A two-sided t test was used to test the differences in quantile normalized pseudobulked data between CM and MM patient samples. Fold changes of genes and pathways were calculated from the mean expression levels in CM and MM from single-cell data. For volcano plot, pseudobulked data was used.

*GSEA*: We performed GSEA using ClusterProfiler[36] on the gene lists ranked by fold changes between MM and CM samples. Gene collections from MsigDB "curated gene sets" were evaluated for enrichment. The most significant upregulated and downregulated pathways (from enrichment scores) were shown.

Figure S1

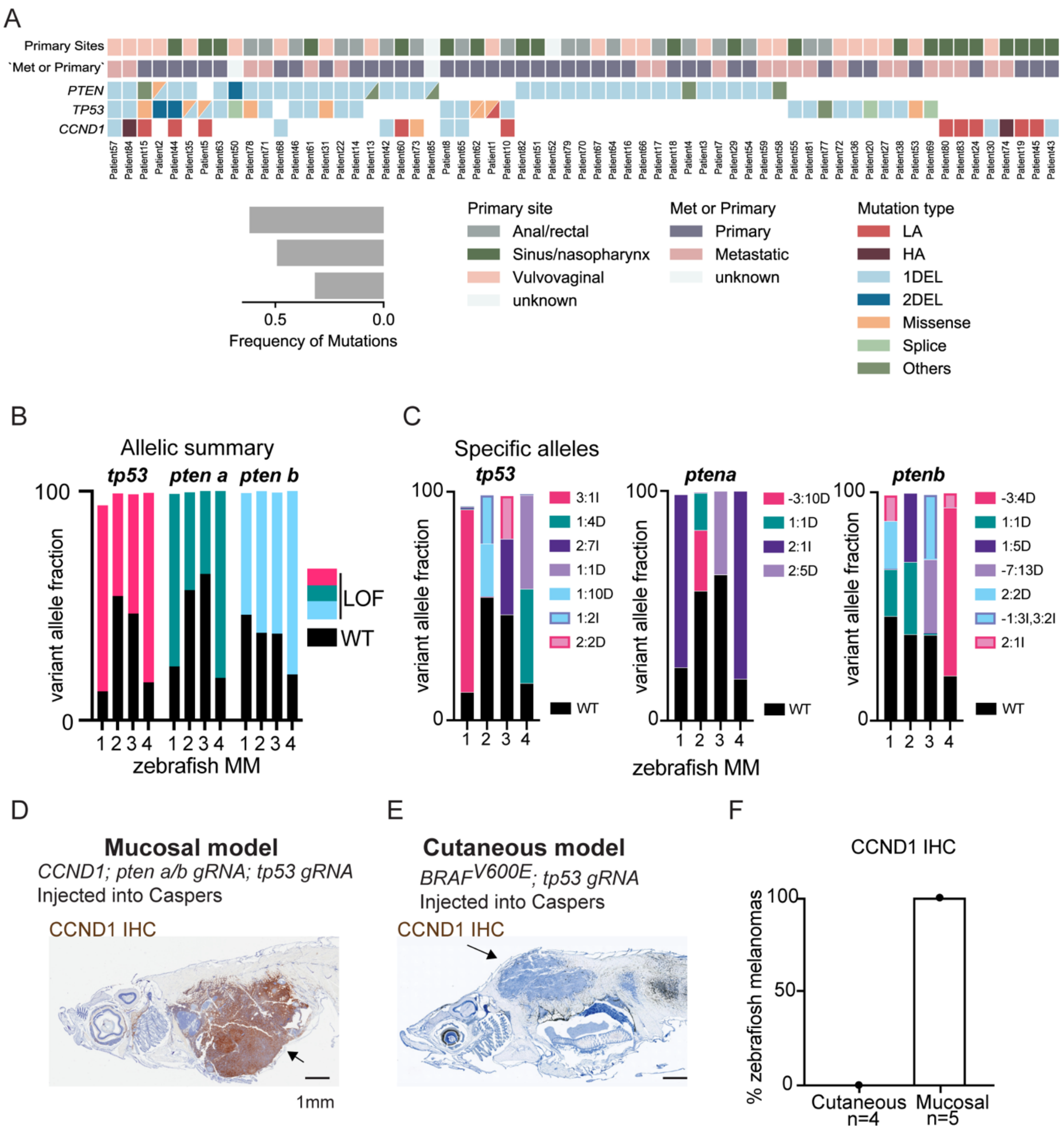


**Fig. S1: Validation of genetic changes in zebrafish MM model.** A) Co-mutation plot from DFCI MM patients with alterations in PTEN, TP53, and CCND1 (n=63/86). LA (low amplification), HA (high amplification), 1DEL (loss of one copy), 2DEL (loss of both copies), Others (indel, nonsense, and truncation). B-C) Variant allelic fraction of B) loss-of-function mutations or C) specific alleles for *tp53, ptena*, or *ptenb* from zebrafish MM. D-F) CCND1 IHC for zebrafish with melanoma for D) MM model, E) CM model, F) % CCND1 positive melanomas quantification.

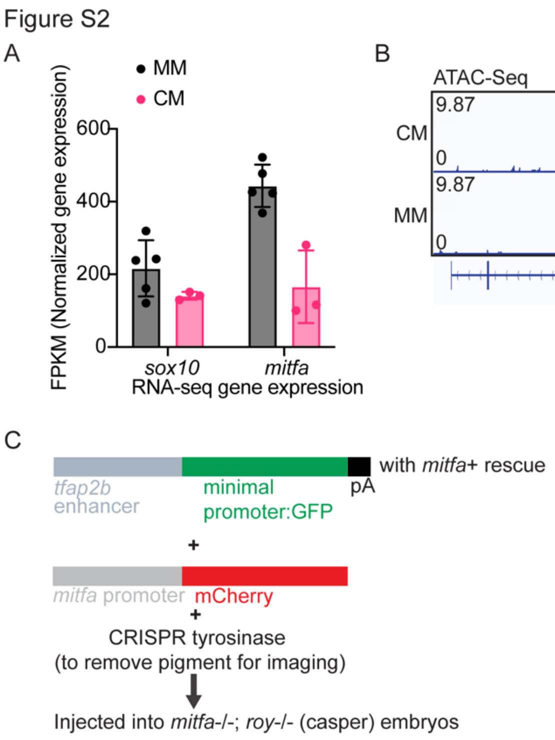


**Fig. S2. Supporting information for the cell state of zebrafish melanoma models.** A) Normalized gene expression of *sox10* or *mitfa* in MM and CM models. B) ATAC-seq tracks illustrating chromatin accessibility at the *pax3b* locus in MM (n=3) vs. CM (n=3). C) Schematic of internal melanocyte reporter experiments.

Figure S3

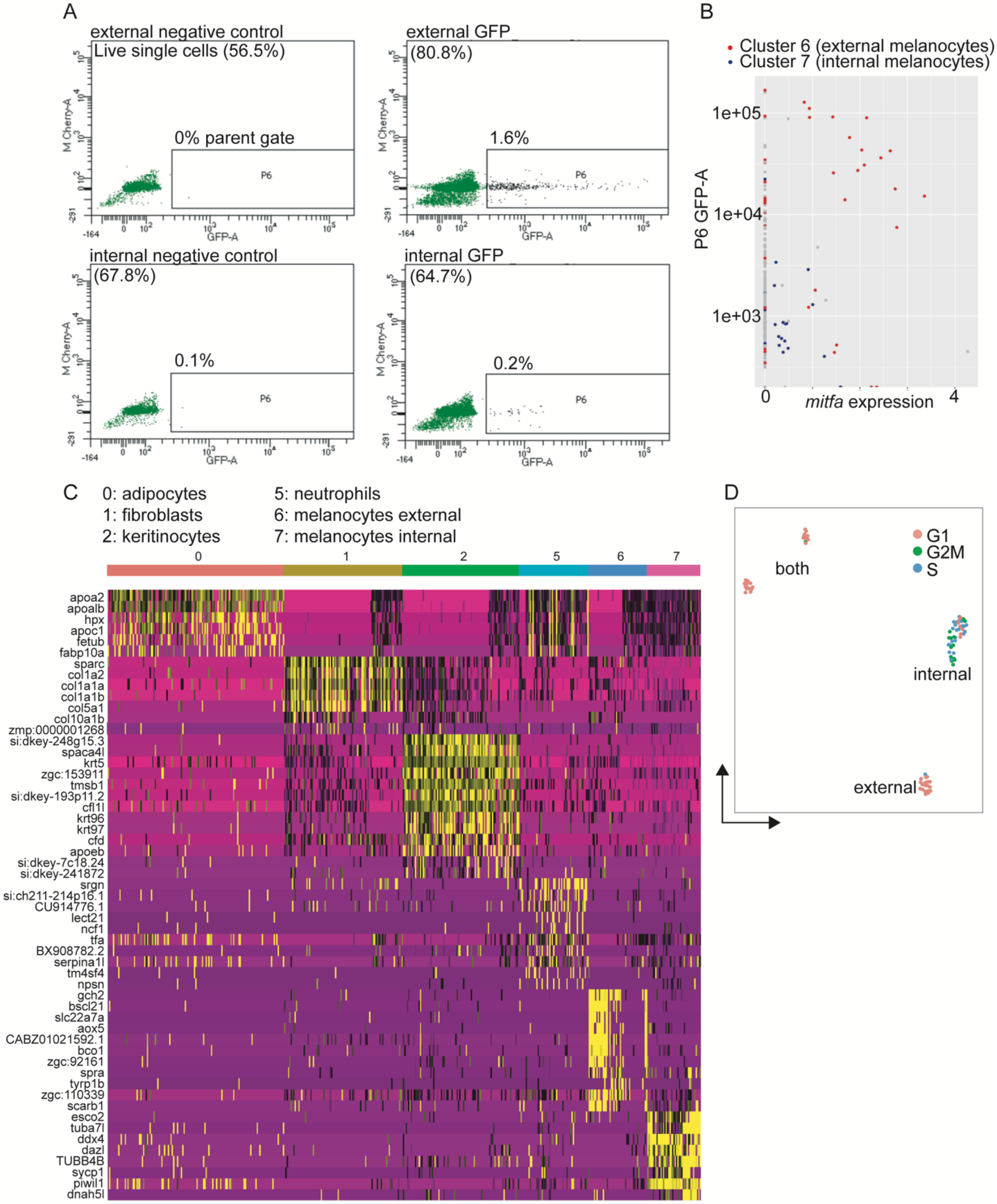


**Fig. S3: Single-cell transcriptomics defines unique populations of adult internal melanocytes in zebrafish.** A) Gating strategy for FACS isolation of GFP-expressing melanocytes from adult zebrafish. B) FACs GFP fluorescence plotted by *mitfa* intensity for all SORT-seq cells. C) Heatmap visualizing single-cell transcriptomes from labeled cell clusters from Fig. 3C. Cluster 3 and 4 were removed from heatmap because they included cells with high mitochondrial gene expression. D) UMAP-plot showing the cell cycle state of internal and external melanocytes from Fig. 3E.

Figure S4

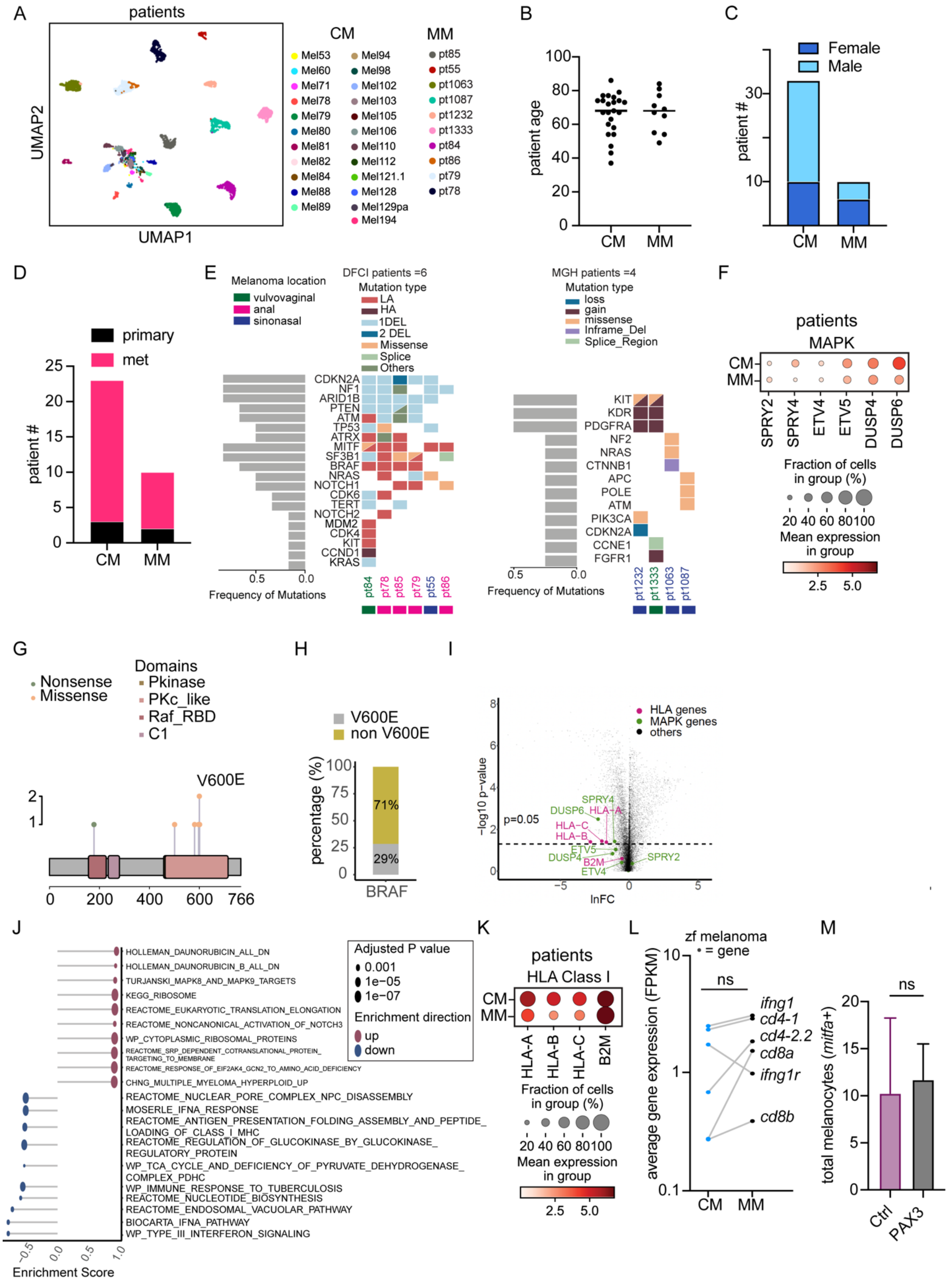
A
patients
UMAP1
UMAP2
CM
MM
Mel53 Mel60 Mel71 Mel78 Mel79 Mel80 Mel81 Mel82 Mel84 Mel88 Mel89
Mel94 Mel98 Mel102 Mel103 Mel105 Mel106 Mel110 Mel112 Mel121.1 Mel128 Mel129pa Mel194
pt85 pt55 pt1063 pt1087 pt1232 pt1333 pt84 pt86 pt79 pt78
B
patient age
CM MM
C
Female
Male
patient #
CM MM
D
primary
met
patient #
CM MM
E
Melanoma location
vulvovaginal
anal
sinonasal
DFCI patients =6
Mutation type
LA
HA
1DEL
2 DEL
Missense
Splice
Others
CDKN2A NF1 ARID1B PTEN ATM TP53 ATRX MITF SF3B1 BRAF NRAS NOTCH1 CDK6 TERT NOTCH2 MDM2 CDK4 KIT CCND1 KRAS
Frequency of Mutations
pt84 pt78 pt85 pt79 pt55 pt86
MGH patients =4
Mutation type
loss
gain
missense
Inframe_Del
Splice_Region
KIT KDR PDGFRA NF2 NRAS CTNNB1 APC POLE ATM PIK3CA CDKN2A CCNE1 FGFR1
Frequency of Mutations
pt1232 pt1333 pt1063 pt1087
F
patients
MAPK
CM
MM
SPRY2 SPRY4 ETV4 ETV5 DUSP4 DUSP6
Fraction of cells in group (%)
20 40 60 80100
Mean expression in group
2.5 5.0
G
Nonsense
Missense
Domains
Pkinase
PKc_like
Raf_RBD
C1
V600E
0 200 400 600 766
H
V600E
non V600E
percentage (%)
71%
29%
BRAF
I
HLA genes
MAPK genes
others
−log10 p-value
p=0.05
SPRY4
DUSP6
HLA-A
HLA-C
HLA-B
ETV5
DUSP4
B2M
ETV4
SPRY2
lnFC
J
HOLLEMAN_DAUNORUBICIN_ALL_DN
HOLLEMAN_DAUNORUBICIN_B_ALL_DN
TURJANSKI_MAPK8_AND_MAPK9_TARGETS
KEGG_RIBOSOME
REACTOME_EUKARYOTIC_TRANSLATION_ELONGATION
REACTOME_NONCANONICAL_ACTIVATION_OF_NOTCH3
WP_CYTOPLASMIC_RIBOSOMAL_PROTEINS
REACTOME_SRP_DEPENDENT_COTRANSLATIONAL_PROTEIN_TARGETING_TO_MEMBRANE
REACTOME_RESPONSE_OF_EIF2AK4_GCN2_TO_AMINO_ACID_DEFICIENCY
CHNG_MULTIPLE_MYELOMA_HYPERPLOID_UP
REACTOME_NUCLEAR_PORE_COMPLEX_NPC_DISASSEMBLY
MOSERLE_IFNA_RESPONSE
REACTOME_ANTIGEN_PRESENTATION_FOLDING_ASSEMBLY_AND_PEPTIDE_LOADING_OF_CLASS_I_MHC
REACTOME_REGULATION_OF_GLUCOKINASE_BY_GLUCOKINASE_REGULATORY_PROTEIN
WP_TCA_CYCLE_AND_DEFICIENCY_OF_PYRUVATE_DEHYDROGENASE_COMPLEX_PDHC
WP_IMMUNE_RESPONSE_TO_TUBERCULOSIS
REACTOME_NUCLEOTIDE_BIOSYNTHESIS
REACTOME_ENDOSOMAL_VACUOLAR_PATHWAY
BIOCARTA_IFNA_PATHWAY
WP_TYPE_III_INTERFERON_SIGNALING
Adjusted P value
0.001
1e−05
1e−07
Enrichment direction
up
down
Enrichment Score
K
patients
HLA Class I
CM
MM
HLA-A HLA-B HLA-C B2M
Fraction of cells in group (%)
20 40 60 80100
Mean expression in group
2.5 5.0
L
zf melanoma
• = gene
ns
ifng1
cd4-1
cd4-2.2
cd8a
ifng1r
cd8b
average gene expression (FPKM)
CM MM
M
ns
total melanocytes (mitfa+)
Ctrl
PAX3

**Fig. S4: Gene expression signatures in MMs.** A) UMAP-plot from scRNA-seq MM and CM patients with individual patients annotated (n=10 mucosal, n=23 cutaneous). B-D) Patient clinical characteristics including B) age, C) sex, and D) whether tumor came from the primary site or from a metastasis. E) Co-mutation plot for scRNA-seq patients showing genetic alterations detected by targeted exon sequencing. For DFCI patients: LA (low amplification), HA (high amplification), 1DEL (loss of one copy), 2DEL (loss of both copies), Others (indel, nonsense, and truncation). F) Dot plot of MAPK target genes in CM vs. MM cells from scRNA-seq. G) Lollipop plot of BRAF mutations. Out of 86 DFCI MM patients evaluated with targeted sequencing, 7 MM patients had protein changes associated mutations in BRAF (nonsense or missense). H) Among the 8% (7/86) MM patients with BRAF mutations, 71.43% (5/7) had non-$BRAF^{V600E}$ mutations, while 28.57% (2/7) had $BRAF^{V600E}$ mutations. I) Volcano plot showing pseudobulked gene expression of MM vs. CM from scRNA-seq. Differentially expressed HLA and MAPK genes are labelled. J) GSEA analysis showing differentially enriched pathways between MM and CM. K) Dot plots of HLA Class I antigen presentation genes in MM vs. CM cells from scRNA-seq. L) Average expression of T-cell genes in zebrafish MM vs. CM. p-value = 0.21, two-tailed Wilcoxon matched-pairs signed rank test. M) Bar graph showing total number of *mitfa*+ melanocytes in control vs. hPAX3 expressing zebrafish embryos. p-value = 0.61, two-tailed t-test.

**Table S1. Zebrafish CRISPR gRNAs from published vectors.**

| Chromosome | start | end | gene | gRNA |
|---|---|---|---|---|
| chr5 | 24087529 | 24087548 | *tp53* | GGTGGGAGAGTGGATGGCTG [7] |
| chr17 | 23673801 | 23673820 | *ptena* | GAATAAGCGGAGGTACCAGG [7] |
| chr12 | 17409879 | 17409898 | *ptenb* | GAGACAGTGCCTATGTTCAG [7] |

**Table S2: Primers used to assess *tp53* and *pten* a/b CRISPR-deletion in zebrafish.**

| | |
|---|---|
| PCR_tp53 Forward | CTGTGTTTGCCAGGAGTACTTG |
| PCR_tp53 Reverse | TATGTGTGTGTATGCGCTTTTG |
| PCR_ptena Forward | GAAGTGTTTTGAACTGCTGT |
| PCR_ptena Reverse | GGAAGTCGTATTGTTACAGCT |
| PCR_ptenb Forward | GCAAGCTCATACCAGGTGTAAA |
| PCR_ptenb Reverse | CCTTCTGAGGAATAAGCTGGAG |

**Table S3: BRAF mutation protein changes in MM targeted sequencing from 86 patients.**

| |
|---|
| p.R178Z |
| p.V600E |
| p.N581Y |
| p.L597S |
| p.E501K |
| p.V600E |
| p.K601E |

**Data S1.** MM mutations from targeted sequencing of 86 patients.
**Data S2:** Clinical and genetic characteristics of MM scRNA-seq patient samples.
